# Clearing-enabled light sheet microscopy as a novel method for three-dimensional mapping of the sensory innervation of the mouse knee

**DOI:** 10.1101/2024.05.28.596316

**Authors:** Frank C. Ko, Spencer Fullam, Hoomin Lee, Shingo Ishihara, Natalie S. Adamczyk, Alia M. Obeidat, Sarah Soorya, Richard J. Miller, Anne-Marie Malfait, Rachel E. Miller

## Abstract

A major barrier that hampers our understanding of the precise anatomic distribution of pain sensing nerves in and around the joint is the limited view obtained from traditional two dimensional (D) histological approaches. Therefore, our objective was to develop a workflow that allows examination of the innervation of the intact mouse knee joint in 3D by employing clearing-enabled light sheet microscopy. We first surveyed existing clearing protocols (SUMIC, PEGASOS, and DISCO) to determine their ability to clear the whole mouse knee joint, and discovered that a DISCO protocol provided the most optimal transparency for light sheet microscopy imaging. We then modified the DISCO protocol to enhance binding and penetration of antibodies used for labeling nerves. Using the pan-neuronal PGP9.5 antibody, our protocol allowed 3D visualization of innervation in and around the mouse knee joint. We then implemented the workflow in mice intra-articularly injected with nerve growth factor (NGF) to determine whether changes in the nerve density can be observed. Both 3D and 2D analytical approaches of the light sheet microscopy images demonstrated quantifiable changes in midjoint nerve density following 4 weeks of NGF injection in the medial but not in the lateral joint compartment. We provide, for the first time, a comprehensive workflow that allows detailed and quantifiable examination of mouse knee joint innervation in 3D.

## Introduction

Joint pain is prevalent in rheumatic and musculoskeletal diseases such as osteoarthritis (OA) and rheumatoid arthritis. In the United States alone, one fourth of the 52 million patients with doctor-diagnosed arthritis have severe joint pain to the point it impairs their daily activities and deteriorates the quality of life^1,2^. While several lifestyle and pharmacological interventions have been implemented or proposed to address joint pain, these treatment strategies are often only moderately efficacious, with limited curative potential. Better understanding of the mechanisms that give rise to joint pain is required for the development of novel therapeutic strategies to improve quality of life for patients with arthritis.

A critical gap in our understanding of the mechanisms underlying joint pain is the lack of a detailed description of the distribution of intra-articular pain-sensing neurons (nociceptors), as well as low threshold mechanoreceptors and proprioceptors. Nociceptors make up > 80% of the sensory innervation of the knee^3^.

Anatomical and immunohistochemical studies have increasingly revealed that the nociceptive innervation of the knee is not static, but rather undergoes profound structural changes with age, joint injury, and OA^4-6^. These observations are predominantly based on two-dimensional histological sections, which preclude precise documentation of nociceptor changes in other parts of the knee joint that are not part of the observed section.

Clearing-enabled light sheet ‘en-bloc’ microscopy is an innovative method that can overcome the limitations of two-dimensional histological approaches. In brief, a tissue of interest is processed through a series of chemicals to render it transparent, which then allows non-destructive optical sectioning by laser light sheet microscopy and reconstruction of the image in 3 dimensions (D)^7,8^. This approach has allowed examination of complete anatomical structures of whole organs comprised of soft tissues such as liver, brain, and intestine^9-11^, but has not yet been reported in whole joints. While we and others have successfully used tissue clearing and light sheet microscopy to visualize 3D features in bone^12-18^, no study to date has rigorously mapped innervation of the whole knee joint, which comprises a mixture of both hard and soft tissues, including bone, articular cartilage, muscle, and synovium. This heterogenous composition of musculoskeletal tissues of the knee joint presents significant challenges when performing clearing-enabled light sheet microscopy, such as refractive index matching, autofluorescence, and antibody labeling due to its extracellular matrix-rich environment.

We have therefore optimized a clearing-enabled light sheet microscopy protocol for visualizing the whole knee joint of a mouse and implemented the method for visualizing and quantifying alterations in nociceptor density following intra-articular injection of nerve growth factor (NGF).

## Materials and Methods

### Mice

Animal studies were approved by the Rush University Medical Center Institutional Animal Care and Use Committee. *C57Bl/6* (JAX664) mice were used, which were group housed 25 mice per cage. Mice were maintained in a pathogen-free facility, subjected to a 12/12 hour light/dark cycle, with *ad libitum* access to standard laboratory rodent chow and water.

### Intra-articular injection of Nerve Growth Factor (NGF)

At 12 weeks of age, male *C57Bl/6* mice were separated into two groups, and saline or NGF (5μl at 100ng/μl, R&D Biosystems, 256-GF-100/CF) were injected into the joint space of the right knee (n = 5 Saline; n = 5 NGF). Left knees did not receive any injections, and the left knees of mice injected with vehicle in the right knee were used as the Uninjected control group (n=5). Mice received intra-articular injections twice per week for 4 weeks. At the end of 4 weeks, mice were perfused with 4% paraformaldehyde under isoflurane anesthesia. Harvested intact knee joints were fixed overnight in 4% paraformaldehyde. Three experimental groups were processed for subsequent analysis: (1) left knee joints from saline injected mice (Uninjected); (2) right knee joints from saline injected mice (Saline); (3) right knee joints from NGF injected mice (NGF).

### Immunolabeling, tissue clearing, and light sheet microscopy imaging

Following fixation, knee joints from all experimental groups were decalcified in 14% ethylenediamine tetraacetic acid (EDTA) for 2 weeks. Decalcified knee joints were then processed for SUMIC, PEGASOS, or DISCO tissue clearing as previously described (Figure 1A)^18-20^. Since we achieved the most transparency with DISCO (Figure 1B), we further modified the protocol (Supplemental File 1) for better binding and penetration of antibodies using 4% SDS/400mM boric acid solution and 0.2% collagenase A (Millipore-Sigma, 11088793001)^21,22^. The samples were then blocked in 6% donkey serum and immunolabeled for protein gene product 9.5 (PGP9.5, 1:500, Millipore-Sigma, SAB4503057-100UG) as a pan-neuronal marker or rabbit-IgG (Abcam, ab37415) as a negative control. Secondary antibody conjugated with AF647 fluorophore (A21245, 1:500; Thermo Fisher Scientific, Waltham, MA, USA) was used to boost the signal from PGP9.5.

**Figure 1.**
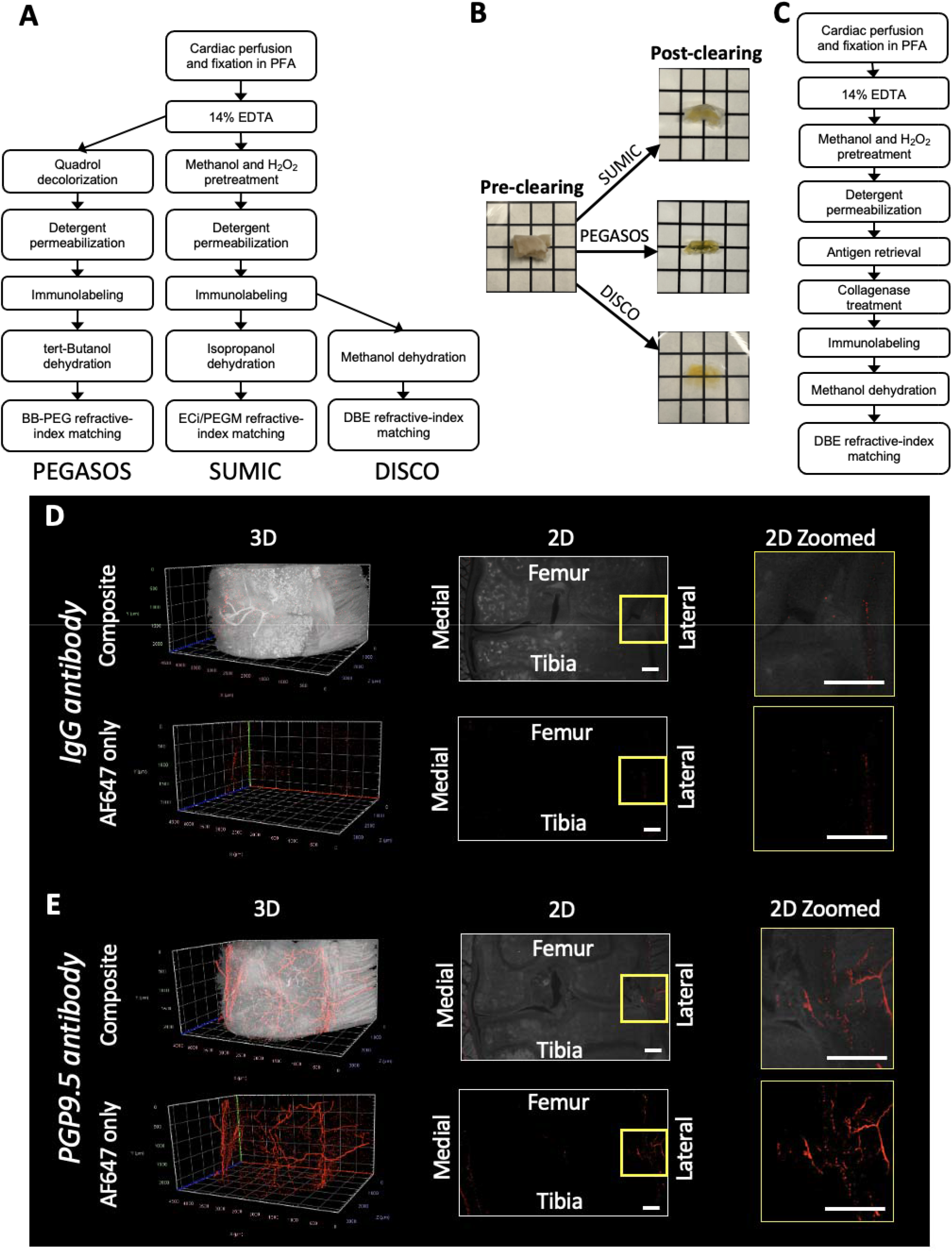
(A) Overview of PEGASOS, SUMIC, and DISCO tissue clearing protocols used for mouse knee joints. (B) Images of mouse knee joints pre-*vs*. post-clearing placed on top of a petri dish with a grid, demonstrating DISCO clearing resulted in complete transparency and ability to see the grid without distortion. Each grid size 8.5 mm x 8.5 mm. (C) Modified DISCO workflow to maximize antibody binding and penetration in and around the knee joint. Light sheet microscopy imaging of the mouse knee joint cleared using modified DISCO procedure stained with (D) IgG antibody (negative control) or (E) PGP9.5 antibody (pan-neuronal). Yellow boxes indicate zoomed in views of the lateral side. 2D slices are imaged at the mid-joint level. Scale bar = 300 μm

Immunolabeled samples were subsequently dehydrated in methanol gradient, delipidated in dichloromethane, and cleared in dibenzyl ether. Cleared knee joints were imaged by ZEISS Lightsheet 7 (ZEISS) to visualize PGP9.5 positive nerves in 3D (Zen Blue 3.7). Samples were oriented so that the light sheet passed in the medial-lateral direction, with the anterior side of the knee facing the objective.

### 3D Image analysis

Light sheet datasets were anonymized, then imported into Imaris 10.1.0 (Oxford Instruments, Abingdon, UK). First, separate ROIs for medial and lateral synovial and capsule tissues at the mid joint were created using autofluorescence contrast in the GFP channel. These ROIs were manually drawn on 4 images, spaced evenly across 67 image planes in the coronal view of the knee, yielding ROIs of 304.18µm in the anterior/posterior axis. At each of the 4 images, the drawn ROI was defined by a straight line from the tibial gutter to the muscle/capsule interface, then a freeform line proximally up the muscle/capsule interface, then a straight line to the femoral gutter, then a freeform line distally down the inner edge of the synovium, connecting the two gutters (Supplemental Figure 2). A 3D ROI was then generated with an Imaris Surface element connecting the manually annotated planes.

**Figure 2.**
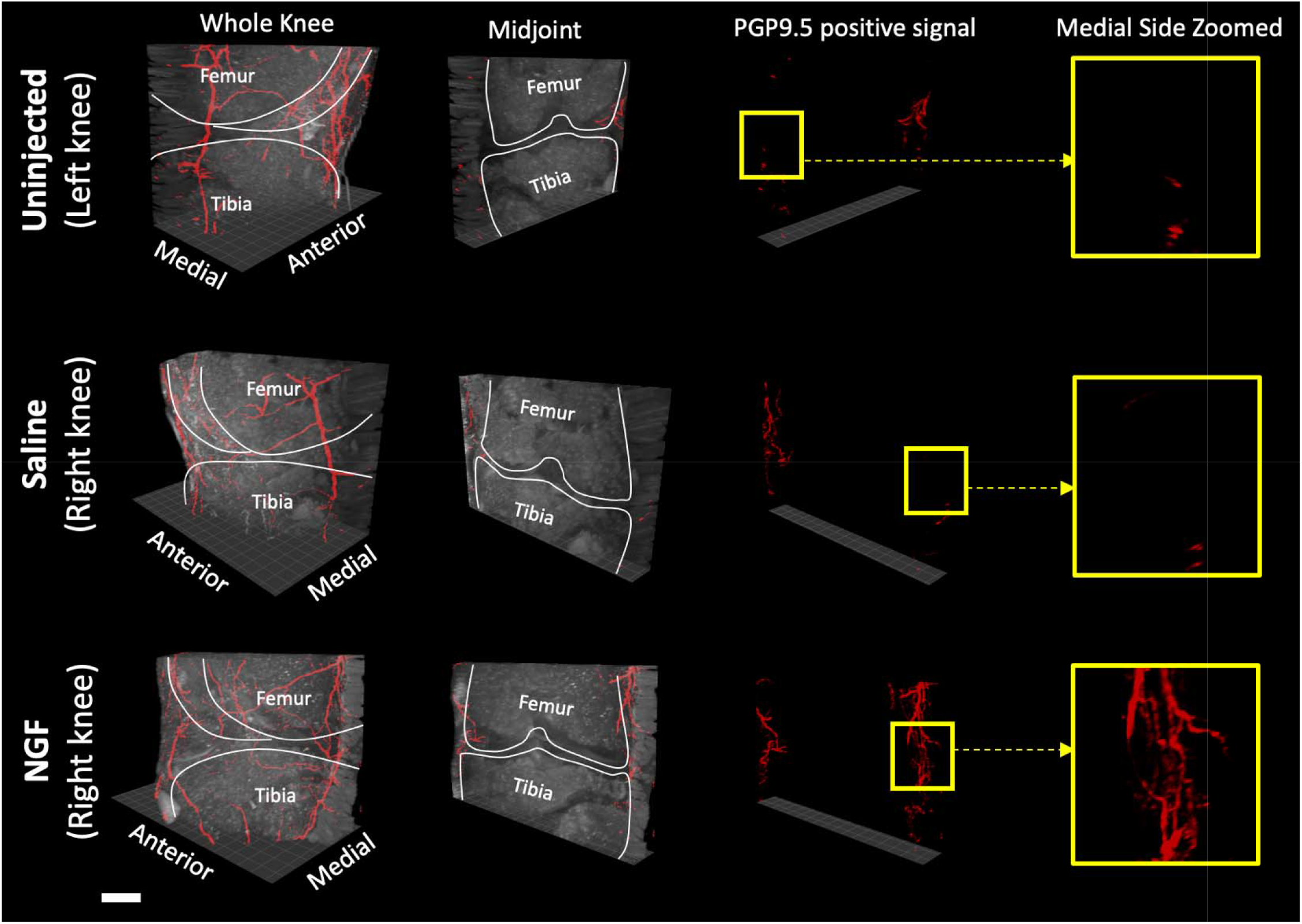
The volume rendering of the knee joint from Uninjected, Saline, or NGF groups. Yellow squares indicate a medial region where NGF group exhibited higher nerve density compared to Uninjected or Saline groups. Scale bar = 500 μm

For each sample, the AF647 channel in the medial and lateral ROIs were separately thresholded, then an Imaris Surface was created to best capture nerve structures while excluding antibody aggregates. Aggregates in medial tissue were filtered out (exclude sphericity > 0.45, Ellipticity (prolate) > 0.4, Ellipsoid axis C Z > 0.95um) slightly differently than lateral tissues (exclude sphericity > 0.45, Ellipticity (prolate) > 0.9, Ellipsoid axis C Z > 0.98um). The volume of the remaining structures was summed, and the volume of the medial and lateral 3D ROIs were also calculated. A volume fraction was calculated for medial and lateral tissues by taking the summed volume of nerve structures and dividing by the volume of the 3D medial or lateral ROI.

### 2D Image analysis

In the previously defined 67 image planes that comprise the lateral and medial ROI, a single plane with the observed greatest positive signal in the AF647 channel was chosen for each side. Fiji was used to outline an ROI in the lateral or medial synovial region on this single plane 2D image as we have reported previously for analysis of traditional immunofluorescence^4^. Thresholding was performed to identify positive signal within the ROI, and the area of positive signal was calculated and normalized to the area of the ROI^23^.

### Statistical analysis

The datasets were unblinded, and the means of volume fraction and area fraction of the uninjected and saline groups were compared to the NGF group with a Dunnett’s one-way ANOVA test (Prism 10.2, GraphPad Software, Boston, USA). Differences were considered significant at *p* < 0.05. Data are reported as mean ± SD.

## Results

### Protocol development

We sought to develop a reproducible protocol that allows clearing of the whole knee joint from a mouse. The main technical challenges that we needed to overcome were 1) poor refractive-index matching with clearing solutions resulting in poor transparency, 2) light scattering and high autofluorescence from the extracellular matrix and heme group, and 3) insufficient penetration of antibodies deep into the tissue. We initially chose to use hydrophobic tissue clearing as it has been shown to result in better transparency compared to aqueous-based tissue clearing approaches^24^. We tested three different hydrophobic-based tissue clearing protocols that have been previously used in skeletal tissues: PEGASOS, SUMIC, and DISCO^18-20^ (Fig. 1A). We found that DISCO achieved full transparency, but PEGASOS and SUMIC resulted in tissues that were still partially opaque (Fig. 1B).

After achieving the full tissue transparency required for light sheet microscopy imaging with the DISCO protocol, we then tested whether peripheral neurons could be labeled and visualized within an intact mouse knee joint by using an antibody against PGP9.5 (a pan-neuronal marker); IgG antibody was used as a negative control. To maximize antibody binding and penetration in and around the knee joint, we modified DISCO protocol to include antigen retrieval by boric acid and extracellular matrix digestion by collagenase (Fig. 1C)^21,22^. The primary antibody signal was further boosted with the secondary antibody conjugated with fluorophores in the far-red spectrum to avoid autofluorescence from dense extracellular matrix. As expected, IgG antibody did not stain any nerves, and very little signal was detected throughout the joint (Fig. 1D). In contrast, PGP9.5-stained samples exhibited clear neuronal staining throughout the intact knee joints (Fig. 1E). In particular, 3D rotational view showed that PGP9.5 positive signals were abundantly expressed on the extraarticular anterior side of the knee (Supplemental Video 1).

To ensure that PGP9.5 signals were observed in the intraarticular space, which is inside the joint capsule (Supplemental Fig. 1A), we examined 2D optical slices within the midjoint region. The mid-joint region was defined by the absence of any ligament attachments to the meniscal horn. We have previously shown, using a conventional histological approach, that dense nociceptor signal in young naïve animals is found in the lateral synovium and insertion of cruciate ligaments^4,23,25^. We similarly found, through our optical slices, that PGP9.5 nerves were abundantly expressed in the lateral synovium as well as cruciate ligament surroundings but minimally in the medial synovium (Supplemental Fig. 1B). This demonstrates that our clearing-enabled light sheet microscopy approach can visualize PGP9.5 positive signals intraarticularly.

### Light sheet microscopy imaging of tissue-cleared knee joints in 3D

Following the protocol development, we sought to examine whether our approach could detect changes in nerve density following NGF injections into the mouse knee joint cavity. We have previously shown using 2D sections that intra-articular injection of NGF led to increased nerve density on the medial side within the mid joint region of the mouse knee^26^. Therefore, we used this NGF injection model in order to test whether we could detect similar changes by tissue clearing and light sheet microscopy imaging. After successfully imaging PGP9.5 signals of the whole knee joint, we created a 3D rendering of the midjoint region to examine if NGF injection alters the nerve density intraarticularly. In the midjoint we qualitatively observed that NGF injection increased the nerve density compared to uninjected or saline injected groups on the medial side (Fig. 2, Supplemental Video 2).

### Analysis of tissue-cleared knee joints in 3D

To confirm our qualitative observations of increased neuronal density in the medial compartment in NGF injected mice, we quantified the nerve density in 3D (Supplemental Fig. 2) at the midjoint. This particular volume was chosen based on our previous studies that demonstrated significant changes in the medial compartment innervation following OA induction^4,23,25^. We found that NGF injection increased the PGP9.5 positive nerve volume fraction by 4.4-fold (*p* = 0.01) compared to Uninjected and by 2.5-fold (*p* = 0.04) compared to Saline injected knee joints (Fig. 3A). The volume and density of PGP9.5 positive nerves on the lateral side did not change significantly following NGF injection (Fig. 3B).

**Figure 3.**
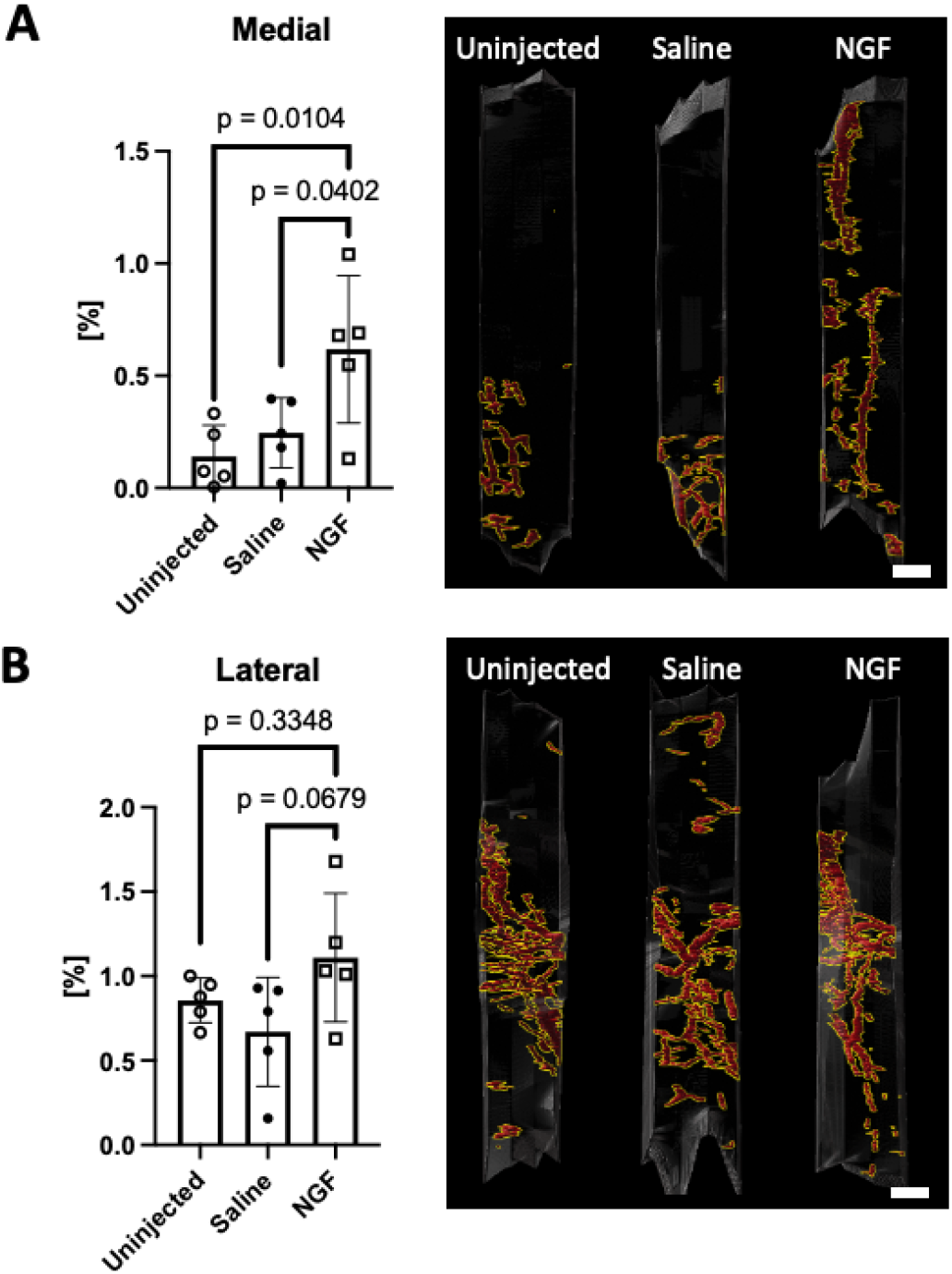
The volume fraction of PGP9.5 positive nerves on the medial (A) and lateral (B) side of mouse midjoint intraarticular space. Scale bar = 100µm

### Analysis of tissue-cleared knee joints in 2D

To accompany our 3D analysis, we performed 2D analysis of optical slice to examine changes in PGP9.5 positive signal following NGF injection. For this study, we used a 2D optical slice at the midjoint with the maximum positive signal from light sheet image datasets (Fig. 4A), and quantification was performed by a blinded observer. Similar to our previous study^26^, we found that NGF injection increased nerve density on the medial side, but not on the lateral side. When we quantified the changes in nerve density, NGF group increased the neuronal density by 3.4-fold (*p* = 0.0022) compared to Uninjected group and 2.2-fold (*p* = 0.012) compared to Saline group (Fig. 4B).

**Figure 4.**
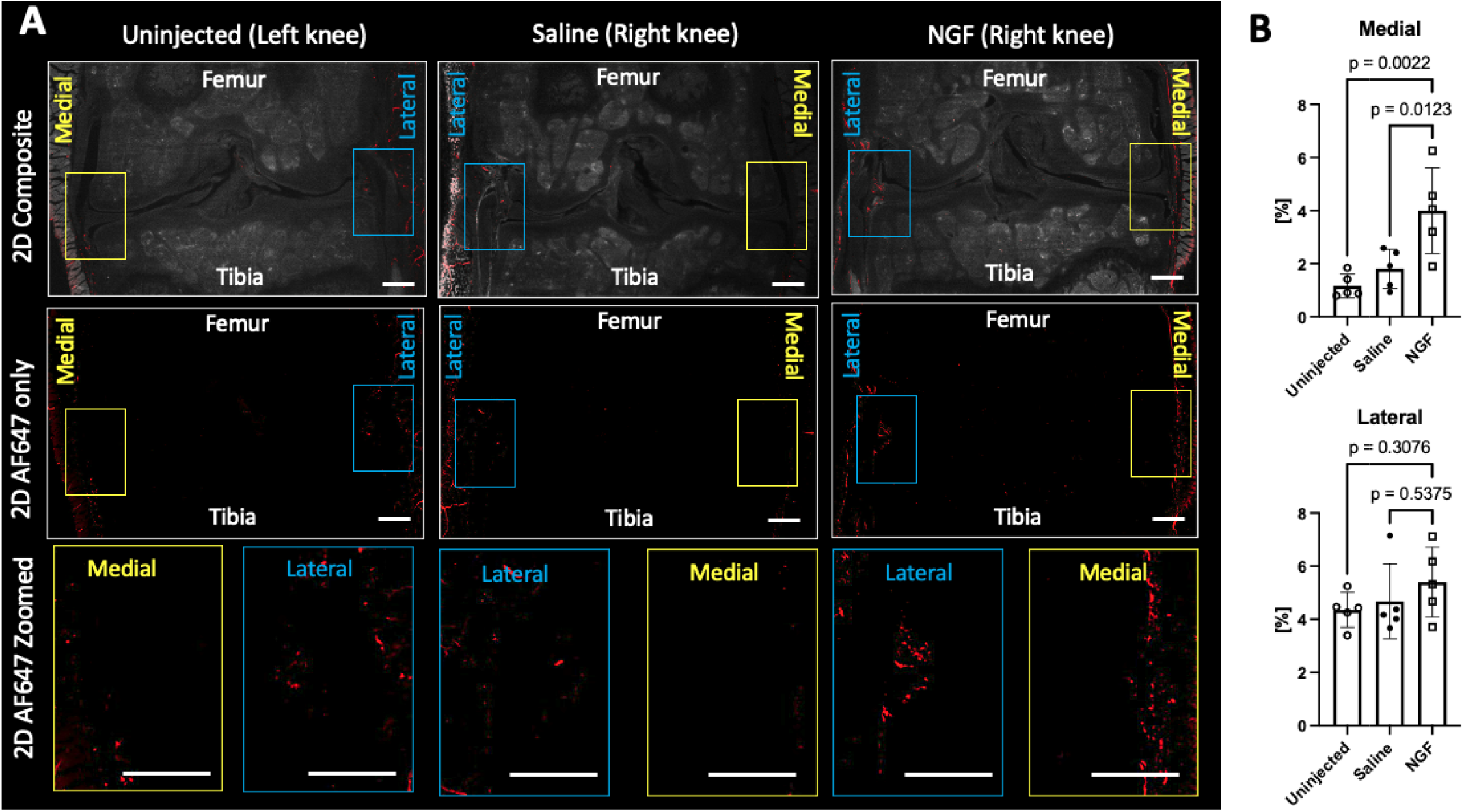
(A) Representative 2D optical sections of the mouse knee joint from each experimental group. (B) The area fraction of PGP9.5 positive nerves on the medial and lateral side of mouse midjoint. Scale bar = 300µm

## Discussion

We developed an experimental workflow that for the first time, allows clearing-enabled light sheet microscopy imaging of an intact mouse knee joint. This was achieved by overcoming several technical challenges associated with examining the combination of hard and soft musculoskeletal tissues in the joint, such as mineralized matrix and dense extracellular matrix proteins. These challenges were addressed by using different enzymes, detergents, antibodies, and fluorophores that allowed labeling and imaging of PGP9.5 positive nerves innervating in the mouse knee. The experimental workflow was able to detect the expected changes in the nerve density following intraarticular NGF injection, suggesting possible applications to other rheumatic disease models that elicit pain responses in mice.

While the hydrophobic-based tissue clearing method quenches endogenous fluorescent proteins, thereby necessitating immunolabeling, this approach allows customization of detecting different targets when using appropriate antibodies. Our study demonstrates that *C57Bl/6* mice with no endogenous fluorescent neuronal reporters can still be used to examine nerves. By using antibodies of different species and fluorophores that do not cross react, our tissue-clearing approach will allow simultaneous imaging and quantifying different neuronal subtypes in and around the knee joint.

In addition to overcoming clearing challenges, we also developed both 3D and 2D analysis methods that enabled quantification of NGF-induced changes in nerve density. While the 2D analysis could be directly translated from methods used for standard immunohistochemical sections, additional filtering steps based on morphology were necessary for 3D analysis to avoid quantification of signal associated with antibody artifacts. The pipeline established here is likely to also be applicable for analysis of vascular changes in the knee joint.

We focused our analysis at the midjoint section to determine changes in the nerve density following intraarticular injection of NGF. This was based on prior studies showing that the medial midjoint synovium of a healthy knee is minimally innervated, but NGF injection significantly enhances the nerve density^4,23,26^. In contrast, the lateral midjoint synovium is highly innervated in the healthy knee and 4 weeks of NGF injection is not sufficient to cause significant changes in the nerve density in that region^4,23,26^. While 3D light sheet images offer opportunities to examine other areas of the knee joint, assessing anterior or posterior nerves will require careful consideration of sample orientation during imaging, such as ensuring the light sheet passes at the anterior-posterior direction for anterior nerves or positioning the posterior side of the knee facing the objective.

Other studies of whole mouse tissue clearing have shown the feasibility of visualizing and quantifying nerves^27-30^, but have never been rigorously examined for innervation in intact mouse knee joints. In addition, these approaches are often time-consuming and require specialized equipment and significant amounts of material that can be cost prohibitive. Our protocol can be performed in a laboratory equipped with standard histological tools, while allowing relatively higher throughput in the number of processed samples with efficient use of reagents. However, several considerations must be made prior to implementing the protocol, such as antibody incubation times or decalcification reagent (eg. formic acid) since knee joints from aged mice or different species will have different extracellular matrix compositions^31,32^ that will likely impact the clearing performance.

In conclusion, we successfully developed and implemented a workflow protocol that allows detailed examination of intact mouse knee joint innervation using clearing-enabled light sheet microscopy. The robustness of our protocol will allow examination of subsets of pain sensing nerves as well as non-neuronal cells that contribute to joint pain. Careful anatomic annotation of pain contributing cells and nerves will pave the way for developing new interventions to treat pain in joint diseases.

## Acknowledgments

Funding for this work was provided by NIH K01AR077679, R01AR060364, R01AR064251, UC2AR082186, R01AR077019, and F31AR083277. The Chicago Center on Musculoskeletal Pain (P30AR079206) provided experimental support. We thank the late Craig Rathwell for his helpful discussions to tissue clearing.

---

**Supplemental File 1**. Detailed protocol template for clearing mouse knee joints

**Supplemental Figure 1.**
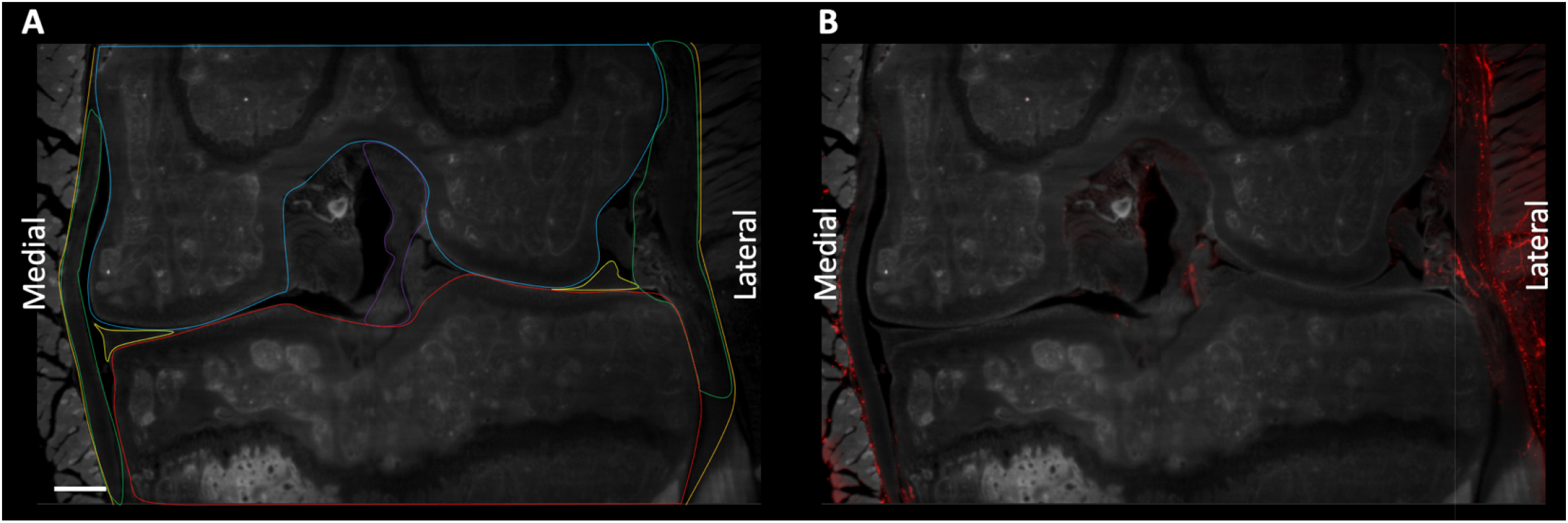
(A) Anatomic features inside a mouse knee joint outlined in different colors. Blue = Femur; Red = Tibia; Orange = Joint capsule; Purple = Anterior cruciate ligament; Yellow = Medial or lateral meniscus; Green = Medial or lateral synovium. (B) 2D optical slice from light sheet microscope imaging of a mouse knee joint stained with PGP9.5. Gray = tissue autofluorescence; Red = PGP9.5 positive signal.

**Supplemental Figure 2.**
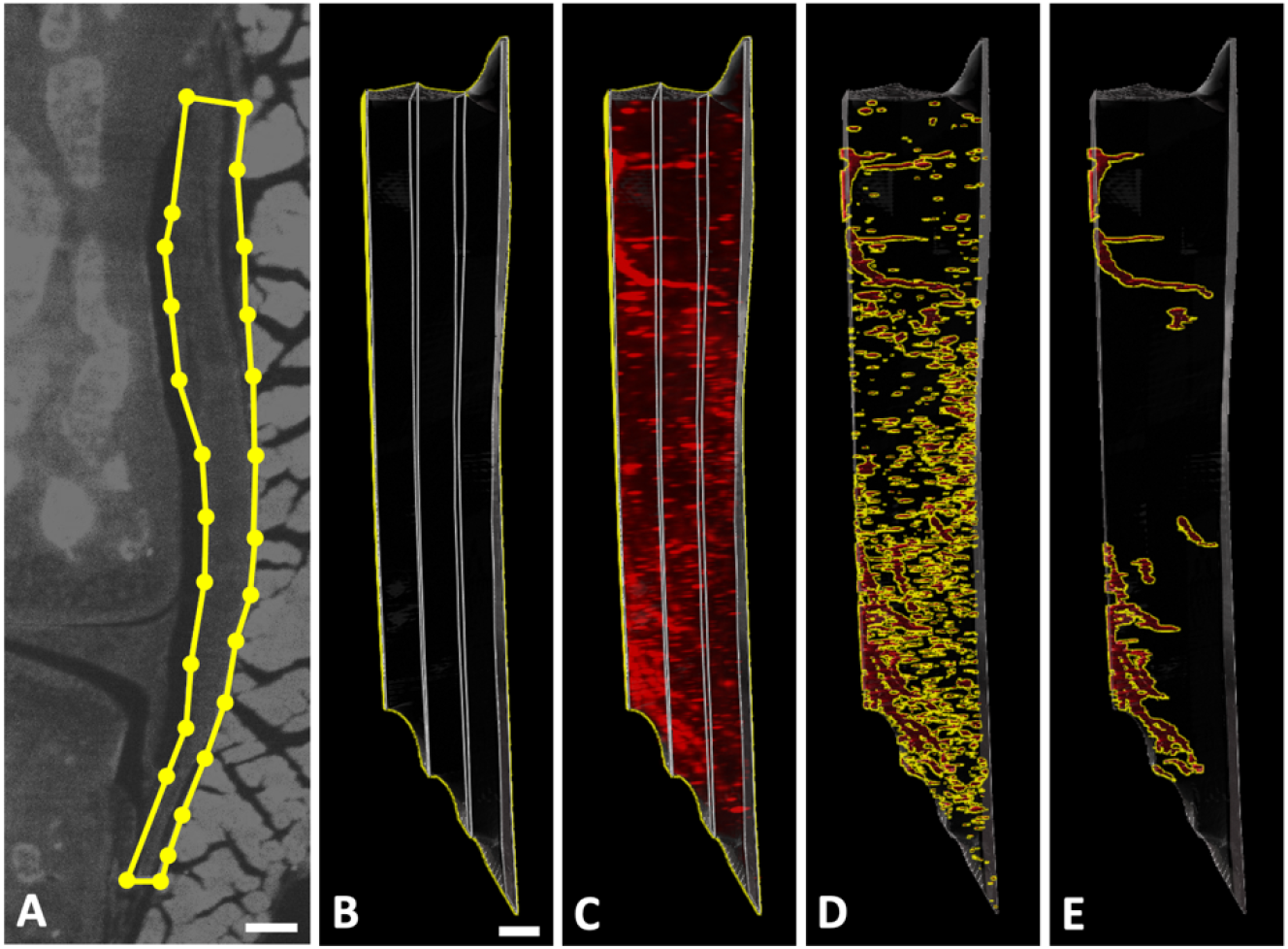
Analytical pipeline for processing light sheet datasets includes (A) manually annotating synovium and capsule tissue in four planes along the mid joint, (B) creating a 3D ROI by interpolating between the four planes, (C) isolating the antibody channel within the ROI, (D) creating Imaris Surface features based on thresholded fluorescence intensity, (E) applying 3 morphological filters to yield the final result. Scale bar indicates 100µm in (A) and approximately 100µm in the 3D projections in (B – E).

**Supplemental Video 1**. Rotating view of PGP9.5 positive signals from the cleared mouse knee joint.

**Supplemental Video 2**. Rotating views of PGP9.5 positive signals from the midjoint region.

