## Supplementary figures and images for "Clearing-enabled light sheet microscopy as a novel method for three-dimensional mapping of the sensory innervation of the mouse knee"

### Supplemental Video 1

#### Slide 1
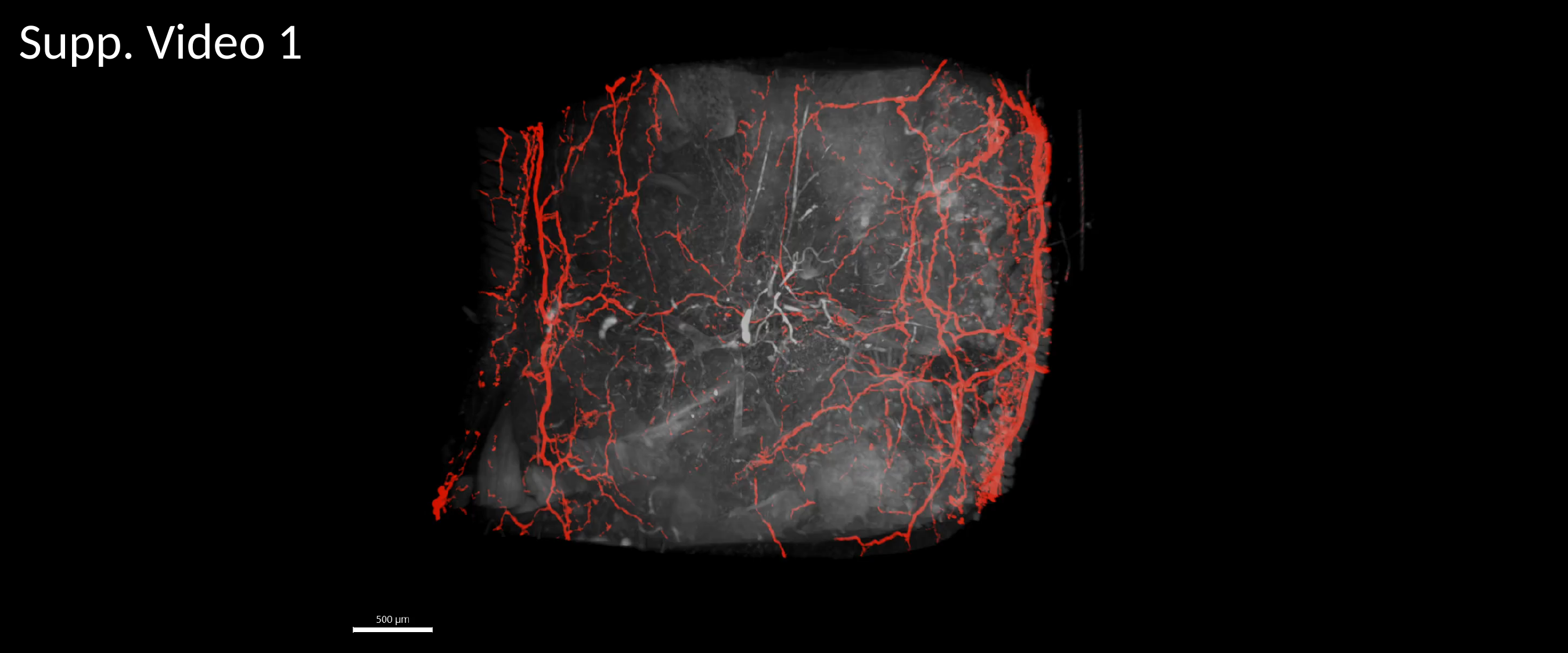

Supp. Video 1

### Supplemental Video 2

#### Slide 1
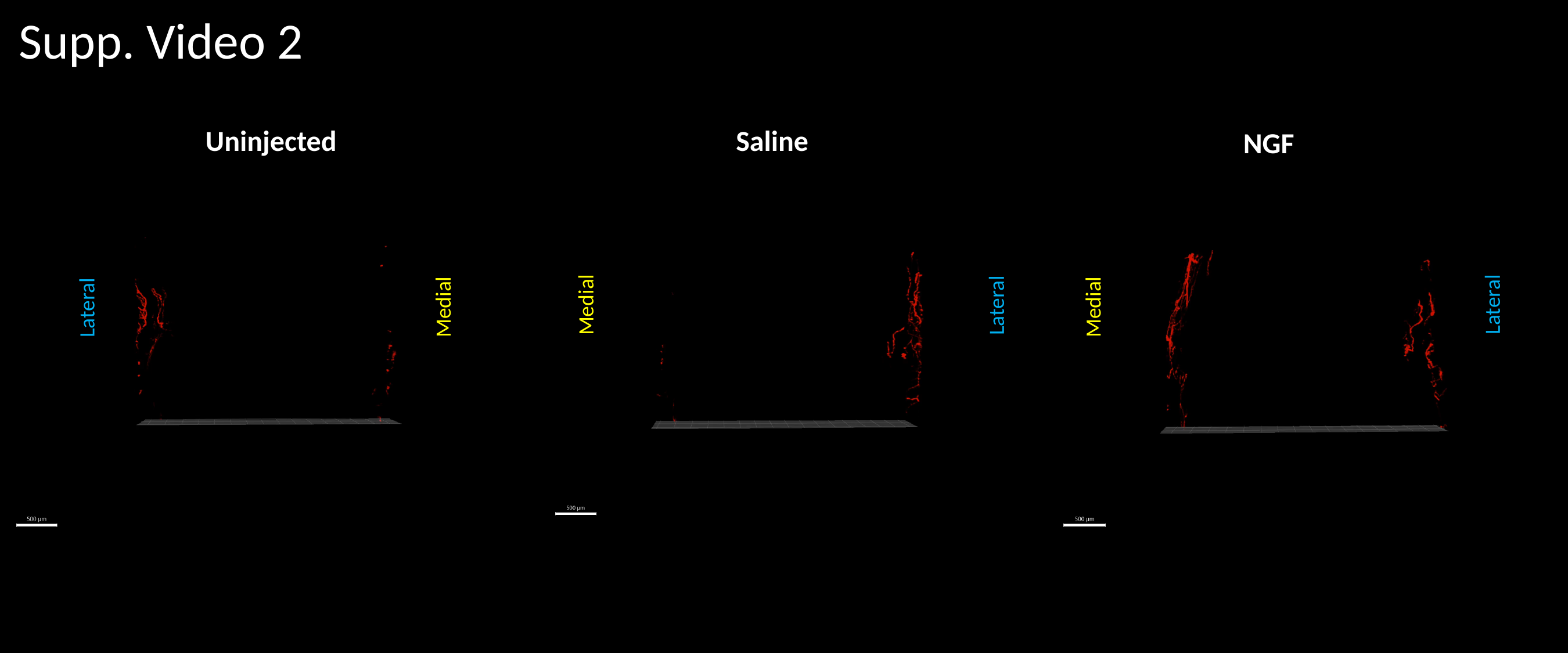

Supp. Video 2
Uninjected
Saline
NGF
Lateral
Medial
Lateral
Medial
Medial
Lateral
